# ESPressoscope: a small and powerful platform for in situ microscopy

**DOI:** 10.1101/2024.01.20.576174

**Authors:** Ethan Li, Vittorio Saggiomo, Wei Ouyang, Manu Prakash, Benedict Diederich

## Abstract

Microscopy is essential for detecting, identifying, analyzing, and measuring small objects. Access to modern microscopy equipment is crucial for scientific research, especially in the biomedical and analytical sciences. However, the high cost of equipment, limited availability of parts, and challenges associated with transporting equipment often limit the accessibility and operational capabilities of these tools, particularly in field sites and other remote or resource-limited settings. Thus, there is a need for affordable and accessible alternatives to traditional microscopy systems. We address this challenge by investigating the feasibility of using a simple microcontroller board not only as a portable and field-ready digital microscope, but furthermore as a versatile platform which can easily be adapted to a variety of imaging applications. By adding a few external components, we demonstrate that a low-cost ESP32 camera board can be used to build an autonomous in situ platform for digital time-lapse imaging of cells. This platform, which we call the ESPressoscope, can be adapted to applications ranging from monitoring incubator cell cultures in the lab to observing ecological phenomena in the sea, and it can be adapted for other techniques such as microfluidics or spectrophotometry. The ESPressoscope achieves a low power consumption and small size, which makes it ideal for field research in environments and applications where microscopy was previously infeasible. Its Wi-Fi connectivity enables integration with external image processing and storage systems, including on cloud platforms when internet access is available. Finally, we present several web browser-based tools to help users operate and manage the ESPressoscope’s software. Our findings demonstrate the potential for low-cost, portable microscopy solutions to enable new and more accessible experiments for biological and analytical applications.

## Introduction

Microscopes are an essential tool in almost all areas of the life sciences. They are used to make analytical measurements, observe dynamic processes at cellular scales, and gain insights into small structures. The availability of consumer and hobbyist technologies like smartphones, miniaturized image sensors, microcontroller development boards, single-board computers, 3D printers, and other rapid prototyping tools and platforms have revolutionized the microscopy field. These tools are approachable and low-cost, making it easier for more people to access and benefit from the latest advancements in microscopy. Innovative laboratories now operate commercial and custom-designed microscopes taking advantage of these technologies, enabling advanced imaging capabilities [1] for scientific exploration and discovery. For example, customized solutions have been developed for super-resolution imaging [2, 3, 4, 5] and malaria diagnostics [6, 7, 8]; while modular platforms have been developed to simplify cutting-edge techniques [4, 9, 10]. These innovations increase the power of microscopy as an analytical tool [11].

Outside of traditional research facilities, the same advances in consumer technologies have enabled affordable, user-friendly microscopes that can easily be customized [12, 13] to meet specific experimental needs [14], and also to make microscopy accessible to a wider range of researchers and students. Many of these microscopes have been designed to promote STEAM education and research in previously underserved communities [15, 16]. An excellent example is the 2 dollar Foldscope [16] and its associated global community for citizen scientists to share what they see in the microcosmos. Other microscopes have been developed with the goal of making specific research techniques more accessible at high performance and low cost. One example is the 3D-printed and open-source OpenFlexure microscope, which is backed by a large user community (estimated 1k users) and is one of the first medically-certified microscopes used to identify malaria in blood smears. Another example is the Planktoscope, a low-cost autonomous stop-flow imaging microscope controlled by a Raspberry Pi to quantify the local biodiversity of plankton using a simple yet powerful fluidic system and microscopy components. These examples show that global impact is achievable for high-performance devices which are made open and affordable, and which develop large user communities. This strategy aims to increase the use of microscopy in life sciences and beyond by providing accessible tools for education, medical diagnosis, and research.

Here, we aim to bridge the gap between the Foldscope and more expensive digital compound microscopes with the development of the ESPressoscope, a low-cost (≤ $10 *-* 30) and simple (build time of ca. 1-3 h) microscopy platform based on consumer-grade microcontroller boards available worldwide. Other components are available off-the-shelf worldwide through online marketplaces such as AliExpress, Alibaba, Amazon, or Taobao. In addition to being compact, easy to source, and easy to build, this versatile microscopy platform requires only a small number of additional hardware components for adaptation into the following five application-specific configurations described in this paper (Figure 1):

**Fig 1.**
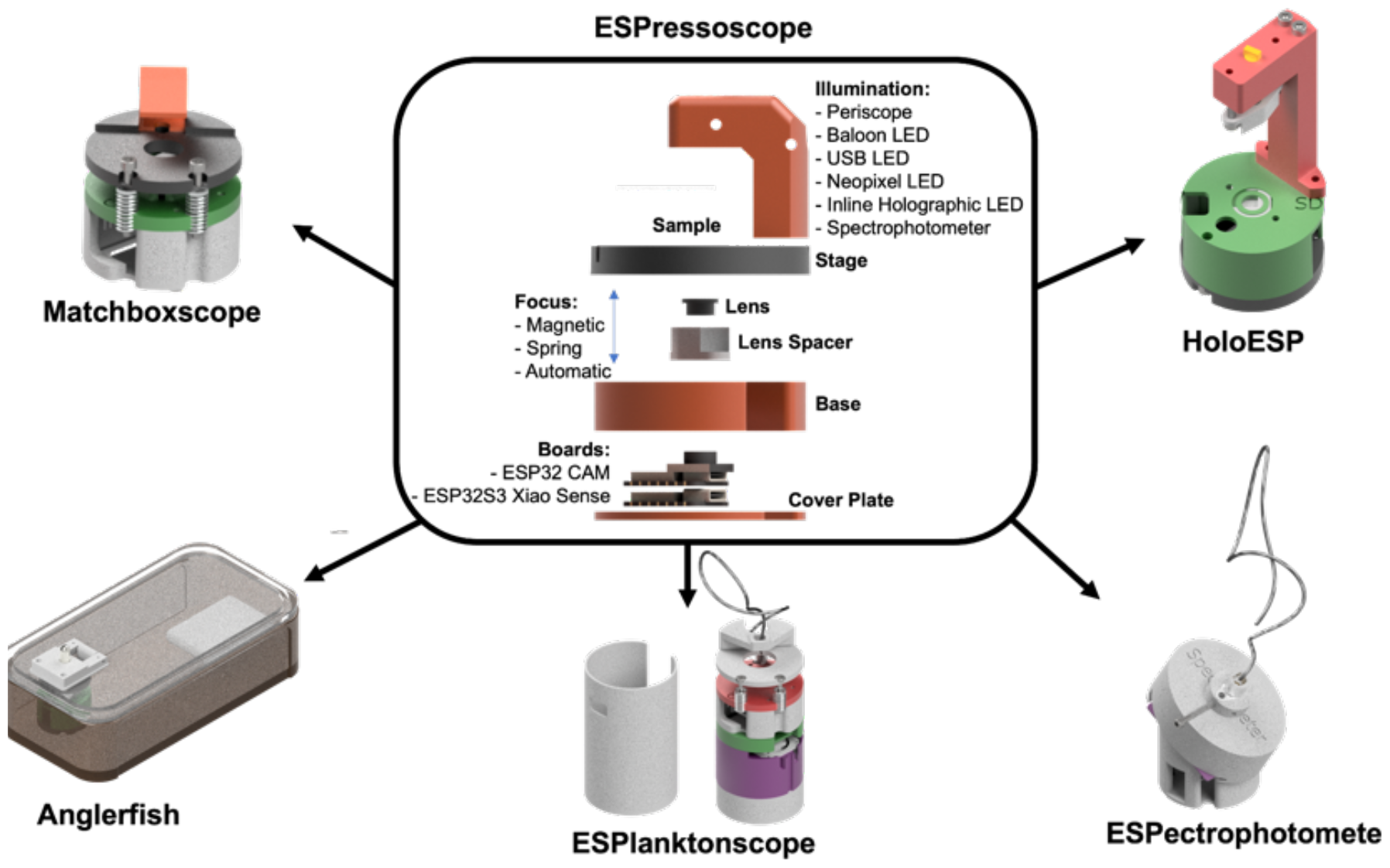
The ESPressoscope platform consists of a microcontroller development board with an integrated camera, 3D printed parts, and a few mechanical parts. From a simple microscope core unit (Matchboxscope), it is possible to fabricate more complex imaging units such as underwater microscopes (Anglerfish), a microscope with an embedded fluidic device (ESPlanktoscope), a spectrophotometer (ESPectrophotometer), and a lensless holographic microscope (HoloESP).

- “Matchboxscope”: A portable brightfield microscope with a resolution between 3 *-* 4 *μm* and a Wi-Fi interface. Multiple modifications are possible to improve illumination, stage movement, z-stacking, etc.
- “Anglerfish”: An underwater microscope for studying aquatic microorganisms underwater
- “ESPlanktoscope”: A simplified version of a flow-imaging microscope with an integrated peristaltic pump
- “ESPectrophotometer”: a compact, digital, low-cost visible-light spectrophotometer
- “HoloESP”: A compact, low-cost, lensless holographic microscope

In all configurations, the camera module can act as Wi-Fi access point for local and remote operation of microscope functionalities, either at the before deployment or throughout the entire deployment. A USB power bank or a lithium-polymer battery can power the board.

An associated website (https://matchboxscope.github.io/) provides detailed documentation on how to set up and troubleshoot ESPressoscope devices, and to enable users to flash the ESPressoscope’s latest firmware directly from a web browser without any further installation of third-party software. This software infrastructure will help anyone to build and operate ESPressoscope-based microscopes.

Our primary objective is to develop a versatile, cost-effective microscopic imaging system which various communities could independently adapt and use, and which would be suitable for in situ deployment in various conditions. To achieve this, we identified the functional requirements for the microscope platform’s design and then defined the essential hardware requirements.

The microscope platform should be capable of imaging various kinds of samples, including samples attached to or mounted on microscope slides, as well as liquid samples with suspended particles, such as in microfluidic devices. Additionally, the microscope platform should be deployable for extended durations in various remote settings with limited connectivity and power sources; extreme deployment scenarios may vary from inside incubators, such as for monitoring cell cultures, to underwater locations. This necessitates support for waterproof enclosures and unattended operation throughout several days and a mechanism to compensate for potential drifts in focus and ambient lighting throughout prolonged experiments. Finally, the microscope should be capable of high-contrast imaging. This requires support for oblique illumination, such as darkfield imaging. We aimed to meet these requirements while minimizing the cost and complexity of the devices by minimizing the overall number of parts required for their construction, and by specifying a hardware platform that could be adapted into configurations specialized for each use-case.

The core module of the ESPressoscope microscopy platform, shared across all ESPressoscope configurations described in this paper, is an Arduino-compatible ESP32 microcontroller board with an integrated camera. A low-cost option is the ESP32-CAM board (Ai-Thinker, China, USD $5) in combination with its ESP32-CAM-MB programming board (China, USD $3). The ESP32-CAM board features an OV2640 CMOS Camera (Omnivision, USA, 1600 × 1200 pixels, 2.2 μm pixel size, Bayer pattern) comparable to image sensors in older smartphones and IP cameras. We have also implemented ESPressoscope configurations with another board featuring the same camera sensor, the XIAO ESP32S3 Sense (Seeed Studio, China, USD $17). Although this board is more expensive, it is smaller and more reliable, and it has an external Wi-Fi antenna factor. Both boards have the option of mounting an SD card for file storage.

The developer board can be turned into a macroscopic imaging device simply by unscrewing its fixed-focus objective lens (*f* #2.2, f’ 4 mm). However, it can be adapted into a high-magnification and high-resolution microscope by adding a few off-the-shelf components (screws, light guide cables, optional electronics) and 3D-printed parts. In designing the system, we prioritized the supply-chain availability of necessary parts, the total cost of the resulting bill of materials, and the ease of independently replicating assembled devices. The resulting microscopy platform achieves a variety of hardware configurations which may offer manual or automatic focus adjustment and transmitted light illumination.

## Results and Discussion

### Matchboxscope: A simple microscope for small places

The simplest configuration, a Matchboxscope with a periscopic illumination mechanism (Figure 2a), can be replicated wherever an ESP32-CAM module and a 3D printer are accessible. Design files, all software, and an in-depth guide to replicating the device are available in the project’s online repository at https://github.com/matchboxscope/matchboxscope. Further documentation is published at https://matchboxscope.github.io.

**Fig 2.**
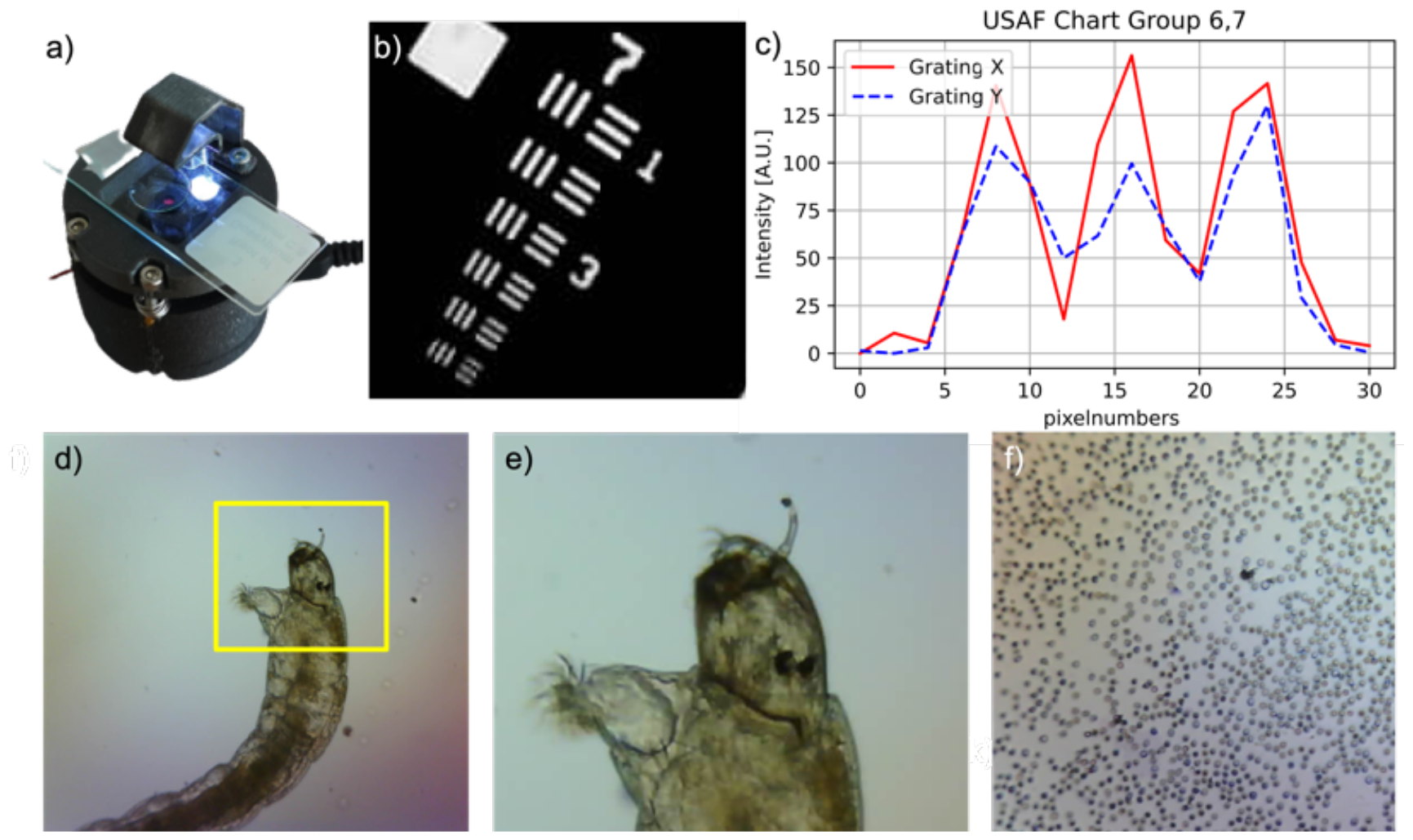
A Matchboxscope configuration featuring periscopic illumination and spring-based focusing mechanism. This simple microscope can resolve features as small as 4-5μm (b,c). Examples of micrographs obtained with the Matchboxscope: A mosquito larvae (d,e) found in a pond and (f) red blood cells.

This initial configuration is a prototypical design suitable as a starting point for more specialized configurations. It can be used for everyday research tasks such as time-lapse imaging in incubators, imaging of microfluidic devices, and portable microscopy for fieldwork. It fits in a pocket (diameter: 52 mm, height: 20-30 mm) and can be powered by an external USB battery. The Matchboxscope’s extreme portability and low cost make it suitable for citizen science projects. For example, one citizen science project sent Matchboxscopes together with simple microfluidic chips [17] to participants for environmental monitoring of microplastics or plankton in water samples.

The simplest optical configuration for a Matchboxscope can be achieved by slightly unscrewing the integrated objective lens of the ESP32-CAM module. Unscrewing the lens moves the static focus from infinity to a finite distance closer to the camera. The magnification is given by

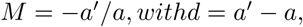

with a the object distance and a’ the image distance, and with a focal length defined by the camera lens 1*/f’* = 1*/a’-* 1*/a*.

An image on the sensor is focused and magnified if *a< a’*. However, to create a compact device with the ESP32-CAM board’s d 4 mm sensor and f’ 4 mm focal length of the objective, we have chosen a’ *≈* 18 *mm* and *≈* 4 *mm*, leading to a total magnification of 4 and an effective pixel size of *≈* 0.6 *μm*. Then the ESP32-CAM board’s f-number of f#2.2 allows a numerical aperture of NA = 1/(2*2.2) 0.23 and thus an optical resolution of *d*(*λ* = 550 *nm*)= *λ/*(2 * *N A*) 1.2 *μm*. Because we use the objective lens in a finite configuration, the effective aperture is smaller, and additional aberration (e.g. defocusing) may further degrade the imaging quality. This configuration maintains the Nyquist sampling criteria if the Bayer pattern is disregarded; otherwise, this configuration is considered to undersample slightly.

Since moving the camera lens would change the effective optical magnification, focusing would be simpler by moving the object relative to the lens to form a sharp image while keeping magnification constant.

### Focus adjustment

For this reason, we developed two possible focus adjustment mechanisms. For manual focus adjustment, the microscopy sample is placed on a 3D-printed spring-loaded or magnet-loaded sample holder which can be moved through the focus by turning one or more of three positioning screws located around the sample. For electronic autofocus, the camera objective lens’ position is linearly adjusted with respect to a fixed camera sensor and sample; the position of the objective lens is actuated using a current-controlled voice coil actuator for M12 CCTV lens (“ML1608” 16X16X8 mm, China). The voice-coil actuator is controlled with pulse-width modulated (PWM) signal from the microcontroller, and it achieves a maximum linear motion of approximately 1 mm discretized with a resolution of 8 bits, leading to an axial step size of approximately 4 *μm*. The electronic autofocus enables simple focus stacking, software-based autofocus to maximize the sample sharpness while varying the focus position, and extended depth-of-field (EDOF) measurements. However, it requires additional electronics and soldering, is vulnerable to slight variation in the magnification factor while focusing, and has a limited focusing range. Supplement Figure 5c shows a focus stack, where a permanent pen on glas was imaged while varying the lens’ position. When electronic autofocus is used, we recommend using it for adjusting fine focus in combination with a manual focusing mechanism for adjusting coarse focus. The Matchboxscope firmware’s contrast-based autofocus algorithm searches the Z position of the objective lens to maximize a relative sharpness metric. For a simple and fast metric, our implementation estimates relative image sharpness by comparing the number of bytes in the compressed JPEG image, since the JPEG compression uses discrete cosine transform (DCT) coefficients in such a way that input images with weaker components at high spatial frequencies will tend to result in an output with fewer bytes; this approach is also used by the OpenFlexure microscope for fast autofocus [18]. More sophisticated sharpness metrics, such as those calculated directly on the distribution of DCT coefficients [19], can be implemented if needed. Supplementary Figure 8 gives an exemplary plot of the sharpness as a function of the lens position.

When images are in focus, the microscope achieves an optical resolution greater than 4 μm, as it resolves the smallest feature group in the 1951 USAF resolution test chart (Figure 2b-c)

### Illumination

The ESP32-CAM module’s integrated LED can be used for sample illumination without additional electrical wiring. In the simplest configuration of the Matchboxscope, light from this LED can be redirected to the sample with a light guide. This can be achieved with a simple 3D-printed periscope whose interior is coated with a reflective surface, such as a coating of waterproof ink from a silver pen marker (Edding 750, Germany), a spray coating of reflective paint, or a layer of reflective aluminum tape, to serve as a light source for transmitted-light microscopy. Alternatively, the periscope can be substituted with a TOSLINK fiber-optic cable normally sold for consumer-grade audio systems (Amazon Basics, 30cm, 4$). In both cases, the redirected light is guided from the ESP32 board to pass from the light guide’s endpoint, through the sample, and into the objective lens; and the endpoint of the light guide defines the illumination mode, which can either be centered, oblique, or darkfield (i.e., outside the objective lens’s NA).

For higher-contrast imaging in critical illumination, we found that an ultra-compact, watertight, coin cell-powered LED light, such as those used in balloon lights, produces a more homogeneous illumination of the field-of-view and is a suitable replacement for the ESP32-CAM module’s integrated LED. Because this light is in a self-contained unit, it can be placed opposite from the objective lens on the other side of the imaging sample, removing the need for a periscopic or fiber-optic light guide. Supplement Figure 3 shows the stability of the light’s intensity over three days. The first trial of a Matchboxscope with this illumination configuration was to detect Schistosoma eggs using a size-selective microfluidic chip (see [17]).

Depending on experimental requirements, more sophisticated illumination methods can also be integrated with the Matchboxscope’s hardware and software. For example, a USB-powered gooseneck lamp from IKEA (USD $4, Sweden), or a NeoPixel LED, LED ring, or LED array (Adafruit, NY, USA, 5-20$) can greatly improve the overall uniformity of illumination or add phase-contrast imaging by dark-field or oblique illumination and is integrated in the device’s firmware. Alternatively, an LED-based system can be substituted with a laser pointer for fluorescence imaging (Figure S5b).

In all configurations, the illuminating end can be moved freely and can be held using 3D-printed components to enable brightfield, oblique, and darkfield configurations. Example images obtained with these four different illumination mechanisms are shown for comparison in Supplement Figure 2.

### Usage

To evaluate the Matchboxscope’s ability to capture long-term timelapse series for in vitro live cell studies, we placed a Matchboxscope unit (spring-loaded manual stage, external LED light source, USB power bank) inside a cell incubator (37 *o C*, 5% CO2, 100% humidity) to capture images every minute over three days to track the growth of a dish of HeLa cells (Supplementary Video 1, Supplement Figure 4). The microscope exhibited imaging drift (200 μm, mostly in the x/y direction) during the first 30 minutes of the timelapse. Thus, the microscope’s mechanical structure requires time to warm up to stabilize before timelapse imaging. At the end of the timelapse, we measured a total drift of about 100 μm within the x/y plane. We hypothesize that this drift is caused by thermal expansion and softening of the Matchboxscope’s 3D-printed thermoplastics, and that the amount of focus drift heavily depends on parameters like materials used, 3D-printing strategy (e.g. infill pattern), and environmental conditions around the Matchboxscope. Supplement Figure 3 shows the lateral drift during a time lapse at room temperature (20*C*, 40% humidity), which is significantly lower and lies at around *±*4 *μm*. Even though the Matchboxscope’s design features an air gap between the sample and the ESP32 camera module, heat buildup in the electronics caused a slight increase in the temperature of the sample-containing petri dish during the cell incubator timelapse experiment. This led to the formation of condensation droplets on the lid which caused non-uniform illumination. This excess heat also induced apoptosis for cells in the dish (Supplement Figure 4); this phenomenon was not observed in an experimental control placed next to the microscope. For this experiment, we had set the Matchboxscope to operate in “continuous” mode (150 *mA* current draw while capturing/streaming images) with an internet connection allowing us to remotely access the microscope’s control interface to monitor the progress of the experiment. The Matchboxscope’s “autonomous” deep sleep mode would be more appropriate, due to its lower power consumption (20 *μA* current draw) and thus lower generation and transfer of heat to the imaging sample between timelapse events. Alternatively, we could have installed a fan to generate airflow between the imaging sample and the Matchboxscope’s camera in order to mitigate heating of the sample from the Matchboxscope.

The Matchboxscope with a magnet-loaded static focus and periscopic illumination configuration was also preliminarily characterized and compared to other microscopes for detecting Schistosoma parasite eggs inside a microfluidic device in [17]. Schistosomatosis is a neglected tropical disease which affects nearly 140 million people annually. [20]. Thus, the ESPressoscope platform can be adapted into a low-cost, portable imaging device for deployment in remote areas to detect Schistosoma parasite eggs in urine samples.

### Anglerfish: A submersible microscope for underwater time-lapse imaging

The study of plankton, biofilms, and other biological dynamics in aquatic surfaces is highly relevant to microbiology, and an in situ imaging tool for studying such dynamics could provide further insights into understanding them. For example, the composition of microbial communities in biofilms can influence processes such as biofouling and even the ecological fate of microplastics [21, 22]. While many researchers have successfully studied biofilm growth under in vitro laboratory conditions, studying their dynamics directly in water has been difficult. Watertight or underwater microscopes in holographic [23], fluorescence [24], or brightfield [25] configurations mostly measure the composition and behaviors of planktonic organisms in fixed volumes rather than measuring microbial colonization processes on submerged surfaces. Furthermore, such microscopes are usually custom-designed, complex, and expensive, with costs as high as - and exceeding - USD $100k.

Here, motivated by open-source and frugal science technologies, we opted for a radically different approach to design Anglerfish, a timelapse-imaging microscope appropriate for recording biofilms, microalgae, plankton, and other aquatic microorganisms in situ, which encloses an ESP32 camera module in a water-tight enclosure. Initially, we upcycled glass fruit preserve jars as water-tight enclosures for the Matchboxscope’s electronics and other water-sensitive components. An alternative approach is to use watertight plastic food containers (e.g. Emsa Clip-and-Close, Germany, USD $4; or Ikea watertight lunchbox “Flottig”, USD $10), which have a less constraining form factor and also can be integrated with a removable microscope slide where biofilm formation is observed (Figure 1). In this case, the microscope slide can be retrieved after the end of in situ time-lapse imaging for post-experiment microscopy with lab instruments at higher magnification and using more sophisticated techniques (e.g. fluorescence microscopy).

### Focus adjustment

The Anglerfish configuration of ESPressoscope features a voice coil-actuated electronic focusing mechanism so that the microscope operator can determine and adjust the focus of the microscope after submerging it. The microscope performs focus bracketing by saving a stack of images while sweeping the focus distance around the desired focus setting at every timelapse event, enabling post-experiment compensation for any small focus drift throughout long timelapse imaging series or, with more complicated image processing, focus stacking.

### Illumination

For sample illumination with Anglerfish, we recommend using a self-contained waterproof battery-driven LED light which outputs light at a high intensity, as it can achieve high imaging contrast despite biofilm growth and requires no additional wiring. This external light module could be substituted with a combination of the ESP32-CAM’s integrated LED and a periscope or TOSLINK cable light guide, but such a substitution will produce weaker illumination and lower imaging contrast. In experiments, we found that the self-contained external LED light can be left on continuously for multiple weeks with one set of batteries. To compensate for the loss of illumination intensity as the battery discharges throughout a deployment, the Anglerfish’s ESPressoscope firmware uses exposure bracketing to acquire and store images on the SD card with a range of exposure parameters for each layer of the focus bracket, enabling selection of images with the best illumination conditions after microscope recovery. This bracketing approach removes the need to rely on automatic white balance, exposure control, and gain control algorithms which could choose inappropriate parameter values in autonomous deployments. This approach enables in situ imaging of diverse biofilm samples without requiring extensive pre-experiment tuning of imaging parameters.

### Power

To maximize power efficiency, the Anglerfish’s ESP32 camera module remains in a low-current deep-sleep mode between timelapse events. Electronic components are powered by a rechargeable lithium-ion battery inside the Anglerfish’s water-tight enclosure. We found that some USB power banks shut down when the current draw falls below some threshold for a given time; such power banks will not power the ESPressoscope properly in “deep sleep” mode, because the ESP32 camera module’s power consumption is too low during deep sleep mode and thus will cause automatic shutdown of the power bank. This problem can be avoided with a DIY power bank in which a lithium-ion battery connected to a 5 V DC step-up converter constantly supplies the ESP32-CAM module with power. The ESP32-CAM can be substituted with a Seeed Studio XIAO camera board, which includes an integrated battery driver so that the lithium-ion battery can instead be connected directly to the board. On average, the ESP32 microcontroller in either board consumes approximately 1000 *mAh* of power per 24-hour period when configured to perform an exposure bracketing series (1/2/5/10/20/50/100/200/500 ms) every minute.

### Improvement with UC2 modules

After setting the focus and initiating time-lapse imaging, the resulting assembly remained operational underwater for at least five days (10,000 mAh power bank, 1 image/minute). The device was functional, but we could not properly capture biofilm formation due to image focusing challenges, even with our focus-bracketing mechanism. Setting up a clear image using the voice coil motor was challenging due to a change in focus upon immersion in water, as water would fill a gap between the lunchbox lid and the glass slide. Further defocusing occurred when transporting the microscope to the underwater deployment site, since small mechanical shocks and vibrations in transit also altered the position of the microscope with respect to the glass surface, resulting in the image being defocused beyond the limits of the voice coil motor’s focusing range. While refocusing was still possible with manual focus adjustment immediately before deployment, the Anglerfish design did not lead to reproducible results in its original implementation, and a larger electronic focusing range was needed.

For these reasons, we modified this version by combining the ESPressoscope approach with modules from the UC2 toolkit of open-source microscopy modules [3]. We formed a finite corrected microscope by combining a Seeed Studio XIAO Sense ESP32 camera board with a customized UC2 cube insert and a motorized z-stage (50mm travel range, NEMA11 stepper motor) holding a 10x 0.25NA finite-corrected objective lens. This assembly fit in the same waterproof container and was successfully deployed in a preliminary test with a 24-hour time-lapse in the Leutra river in Jena, Germany (Figure 3c). In this test, the microscope was operated without the power-saving deep-sleep mode, resulting in high power consumption from the Wi-Fi electronics throughout the experiment.

**Fig 3.**
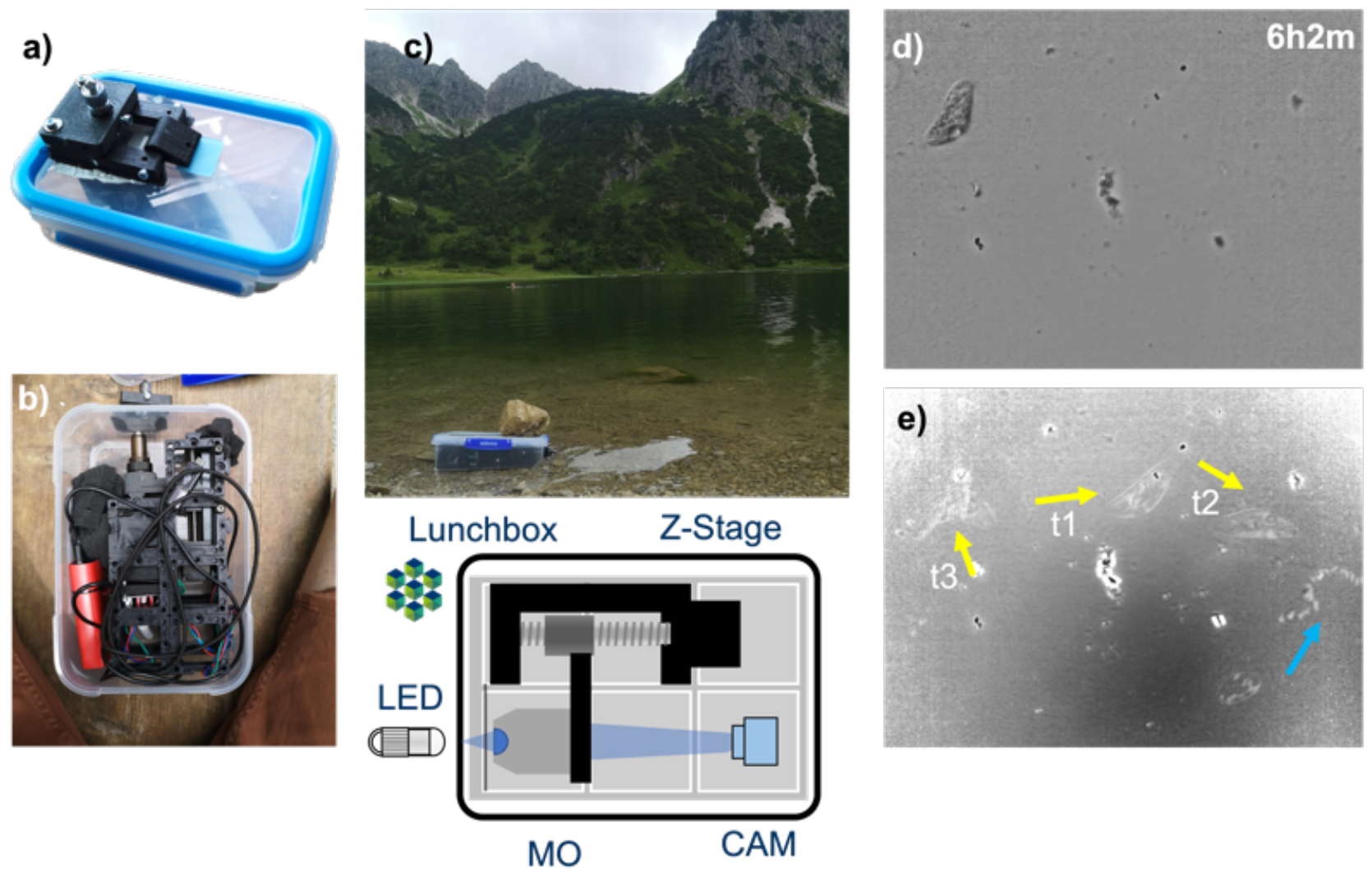
The Anglerfish: a submersible and automatic microscope for underwater operation, built using (a) a Matchboxscope with additional 3D-printed components, and (b) by combining the ESPressoscope software on an ESP32 camera module together with modules from the UC2 microscopy toolkit. With deployment in a pond (c) and application of flat-field correction to acquired images, the microscope allows observation of initial biofilm formation and microbial behaviors, including (d) a Paramecium crossing the field-of-view. (e) A variance projection of the timelapse visualizes the movement of different particles over time, with yellow arrows indicating the movement of the Paramecium at different timepoints and the blue arrow indicating the movement of a smaller microorganism.

Using the ESPressoscope’s web-based GUI, we achieved precise focus of the objective relative to the Anglerfish’s mounted glass slide so that its water-facing surface was in focus. The larger focusing range with the motorized z-stage simplified the focusing of the sample, and the higher-quality objective lens significantly improved image quality and ease-of-use in our test. The ESPressoscope’s autofocusing algorithm successfully compensated for temperature-induced focus drift resulting from the day-night cycle.

The 24-hour timelapse was acquired with flat-field correction of the image sensor performed in Fiji after downloading all images to a local computer [26] in order correct for illumination inhomogeneities. During the experiment, several organisms were observed crossing the field of view (Figure 3d-e). Especially after 20 h, many small organisms were recorded on the glass surface (Supplementary Video 2). However, the microscope’s resolution was insufficient to identify smaller details. One reason for this is that the lunchbox contributes to optical aberration and scattering, which reduces the maximum possible resolution. One solution would be to drill a hole in the lunchbox and seal it with a glass slide, at the risk of compromising watertightness. An alternative solution could be to replace the lunchbox with a more expensive IP66 electric enclosure with a clear window.

Our results with the Anglerfish configuration demonstrate the ability for ESPressoscope configurations to be upgraded for improved imaging performance depending on operational requirements, including through combination with modules from other microscopy hardware toolkits.

### ESPlanktoscope

Thanks to the layered hardware architecture of the ESPressoscope, additional modules can easily be stacked on top of each other to integrate additional functions. To this end, we prototyped an ESPlanktoscope configuration (Figure 4) by adding a small 3D-printed peristaltic pump derived from an open-source hardware design [27]. Like in the original Planktoscope design [28], the flow rate of a liquid sample across the ESPlanktoscope’s field-of-view can be controlled to image individual microscopic particles suspended in a large volume of input sample.

**Fig 4.**
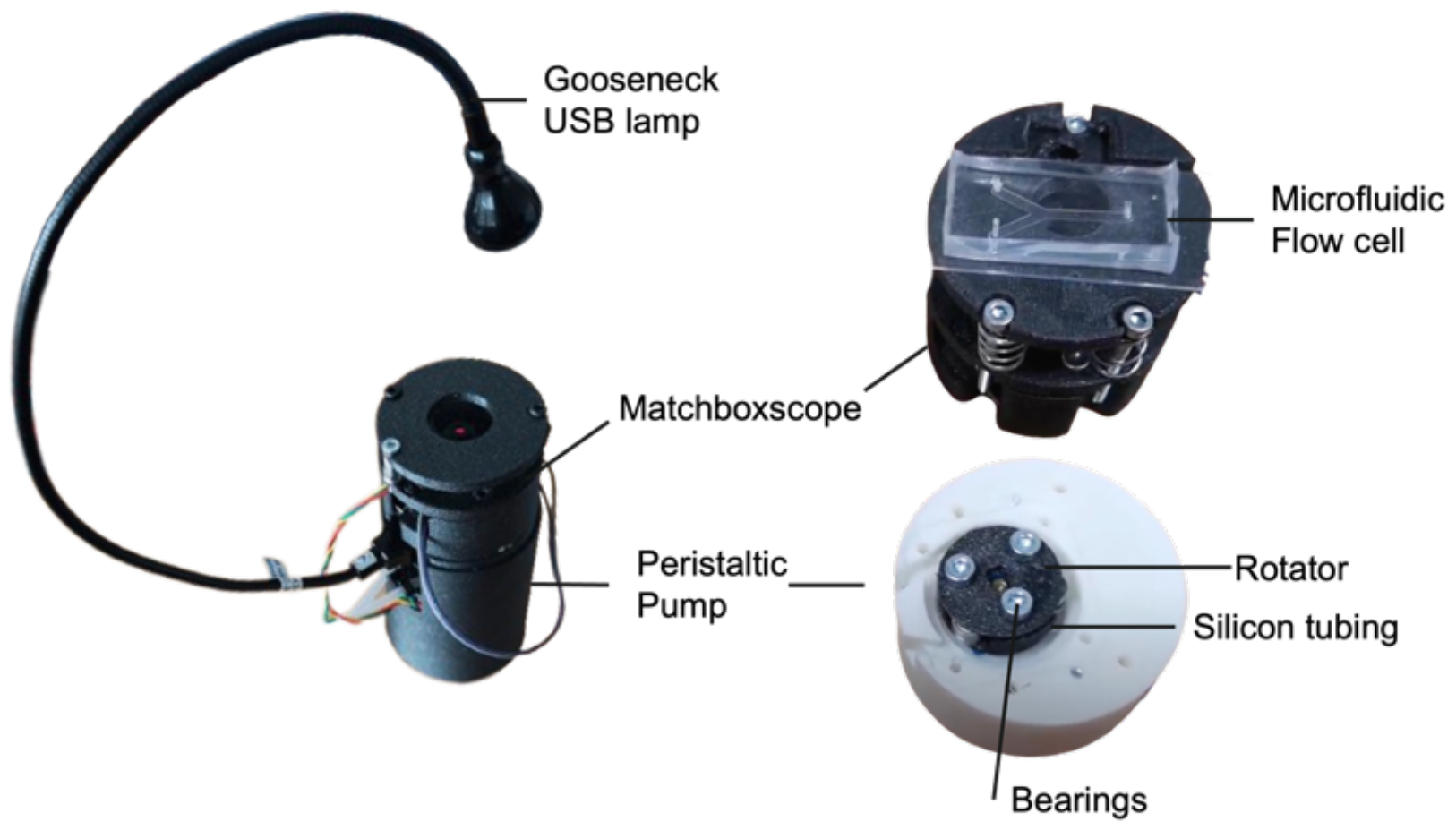
The ESPlanktoscope combines the standard Matchboxscope (top) with an additional ESP32 board for driving a peristaltic pump. Being a Matchboxscope extension, all possible Matchboxscope modifications explained before (different light sources and focusing capabilities) can also be applied to the ESPlanktoscope.

The ESPlanktoscope’s 3D-printed peristaltic pump enables the microscope to continuously measure samples at a high rate (approx. 500 μL/min). To this end, the flow can be controlled in positive and negative directions to move liquid back and forth. Ball bearings reduce the frictional resistance between the rotor and tubing during the pumping process so that the torque provided by the pump motor is sufficient to pump larger samples at a flow rate through the field of view of the microfluidic flow cell.

The ESPlanktoscope’s peristaltic pump is actuated with a stepper motor (28byj-48, 5$, China) which is controlled by a ULN2003 (Texas Instruments, Texas, USA) motor driver. For an ESPlanktoscope using the ESP32-CAM module, which does not provide enough GPIO pins to control the motor directly, the motor driver must be controlled by a second ESP32 microcontroller board connected to the ESP32 camera module, as the ESP32 camera module does not offer enough GPIO pins to control the motor directly; in such a configuration, the ESP32 camera module sends a PWM signal to the second microcontroller in order to set the flow rate of the pump. By contrast, the Seeed Studio XIAO camera board provides enough GPIO pins to control the motor driver directly without the requirement for a second microcontroller board.

Figure 4 shows an assembled ESPlanktoscope unit equipped with an IKEA lamp which can be used for brightfield, oblique, and darkfield illumination. This gooseneck lamp provided a high degree of flexibility to align the light source relative to the homemade microfluidic chip and relative to the tubing connecting the microfluidic chip to the pump. A video showing the device in operation can be found in Supplementary Video 3.

Our prototype ESPlanktoscope configuration demonstrates the feasibility of combining the core parts of the ESPressoscope platform with additional hardware modules to provide new imaging capabilities.

### ESPectrophotometer

Many different open-source designs exist for making compact digital spectrophotometers by combining a camera with a reflection grating (e.g. a CD/DVD) or a transmission grating [29]. This inspired us to extend the functionality of the ESPressoscope into a digital spectrophotometer, the ESPectrophotometer.

Our ESPectrophotometer prototype is implemented by adapting a low-cost transmission grating (Diffraction Grating Slide-Linear 1000 Lines/mm, 1$) with a tilt angle of 35° in front of the camera. The entrance slit for light consists of an FDM 3D printer hotend nozzle (0.1 mm, eBay, 5$) placed approximately 40 mm from the grating. This off-the-shelf brass-machined part guarantees a much higher precision compared to 3D-printed designs. The 3-D printer nozzle creates an angular independent point source of light which passes through the tilted diffraction grating. Only the first diffraction order enters the camera module, which is equipped with an objective lens to focus the spectrum on the sensor.

Software running in a web browser receives camera frames from the ESP32 camera module using the WebSerial browser API, while open-source client-side Javascript software adapted from [**Gaudilab2023npm**] directly processes and visualizes the spectrophotometry data. In this case, the ESP32 camera module is connected to the computer using a USB cable rather than Wi-Fi. The software is provided as part of ESPressoscope’s freely-accessible documentation bundle (https://matchboxscope.github.io/spectrophotometer/espectrophotometer.html) and does not require installation. The software allows the user to select the position of the spectral components before a 1-D spectrum is displayed as a line plot. In Figure 5c-e, we measured the spectrum of a cellphone’s flashlight. The spectrum exhibits a clear dip in the green region of the white light spectrum as is typical for white LEDs. We have not characterized spectral resolution and accuracy in our preliminary demonstration of the feasibility of the ESPectrophotometer as a realizable adaptation of the ESPressoscope platform.

**Fig 5.**
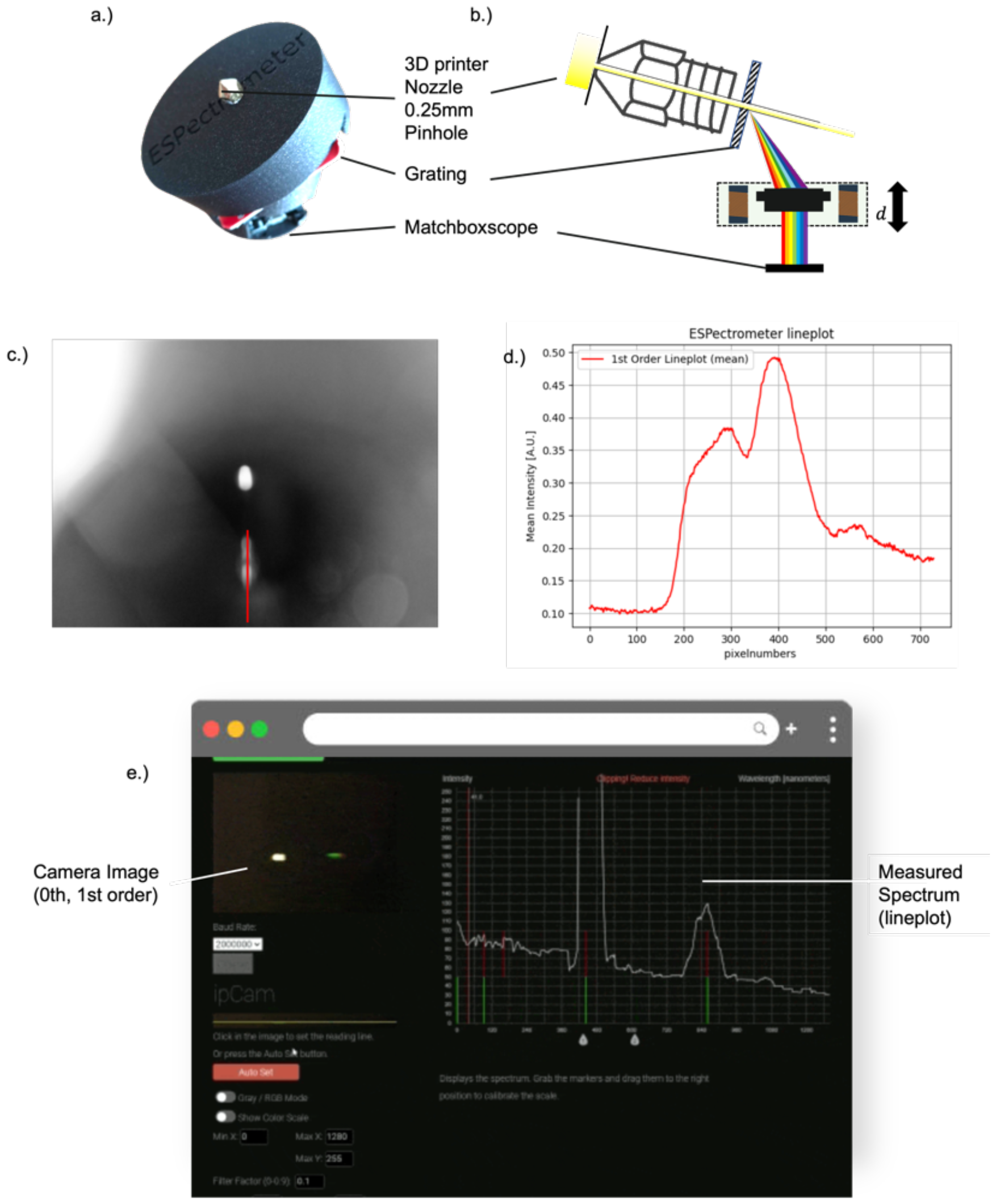
Converting the ESPressoscope into a spectrophotometer requires the objective lens to be mounted on the camera and a 3D printing nozzle to be added as a pinhole. b) An adjacent grating diffracts the light before it gets focused on the sensor. c) The resulting image on the camera sensor can be converted into a spectrum by drawing a d.) line plot through the intensity signal along the first diffraction order. e) A browser-based tool derived from the [30] receives frames from the ESP32 camera and draws the lineplot. For quantitative pixel/wavelength measurements, the ESPectrophotometer would require calibration.

Our prototype ESPectrophotometer configuration demonstrates the feasibility of adapting the ESPressoscope into configurations which integrate real-time image processing and visualization, as well as the versatility of the ESPressoscope platform for use beyond the life sciences.

### HoloESP

Lensless imaging provides the ability to capture in 2.5D and thus refocus an acquired image with digital post-processing. By contrast, the Matchboxscope’s fixed-focus setting poses challenges for dynamic imaging. This limitation can be solved by inline holographic imaging as implemented by the HoloESP configuration of the ESPressoscope platform. A simple narrow-band LED (450nm +/-20nm) is mounted approximately 80 mm from the sample and spatially filtered with an FDM 3D printer hot-end nozzle (0.1 mm). This creates a point source of light which illuminates the sample with weakly coherent spherical waves, producing a very simple and inexpensive inline holographic microscope. The captured holograms are numerically backpropagated using a custom PyScript module which can run in the web browser of a computer connected to the HoloESP microscope. This algorithm numerically refocuses the raw hologram and outputs a real-time display of its back-propagated result. To image small particles such as dust particles, we removed the black plastic cap holding the threaded objective lens provided with the ESP32-CAM module’s camera sensor and instead applied the sample droplet directly to a coverslip placed directly on the sensor window. According to the Nyquist criterion, the approximate resolution should be about twice the pixel size (3 μm). Figure 6 shows the basic working principle, and some numerically refocused microplastic particles that were found in shower gel; the backpropagation algorithm successfully refocused the raw image into the sample plane and formed a sharp image of the microparticles (Figure 6 c-e). The HoloESP achieves a resolution of approximately 3 μm measured by a line plot at the smallest visible structures. Imaging of E. coli bacteria suspended in a liquid sample produced images with very little contrast, resulting in a holographic reconstruction with a very low signal-to-noise ratio. Over time, E. coli sunk down and started attaching to the glass surface; this process was captured by the HoloESP’s camera. However, due to the low signal-to-noise-ratio (SNR) of acquired data, it was impossible to reconstruct the holograms to identify individual bacteria.

**Fig 6.**
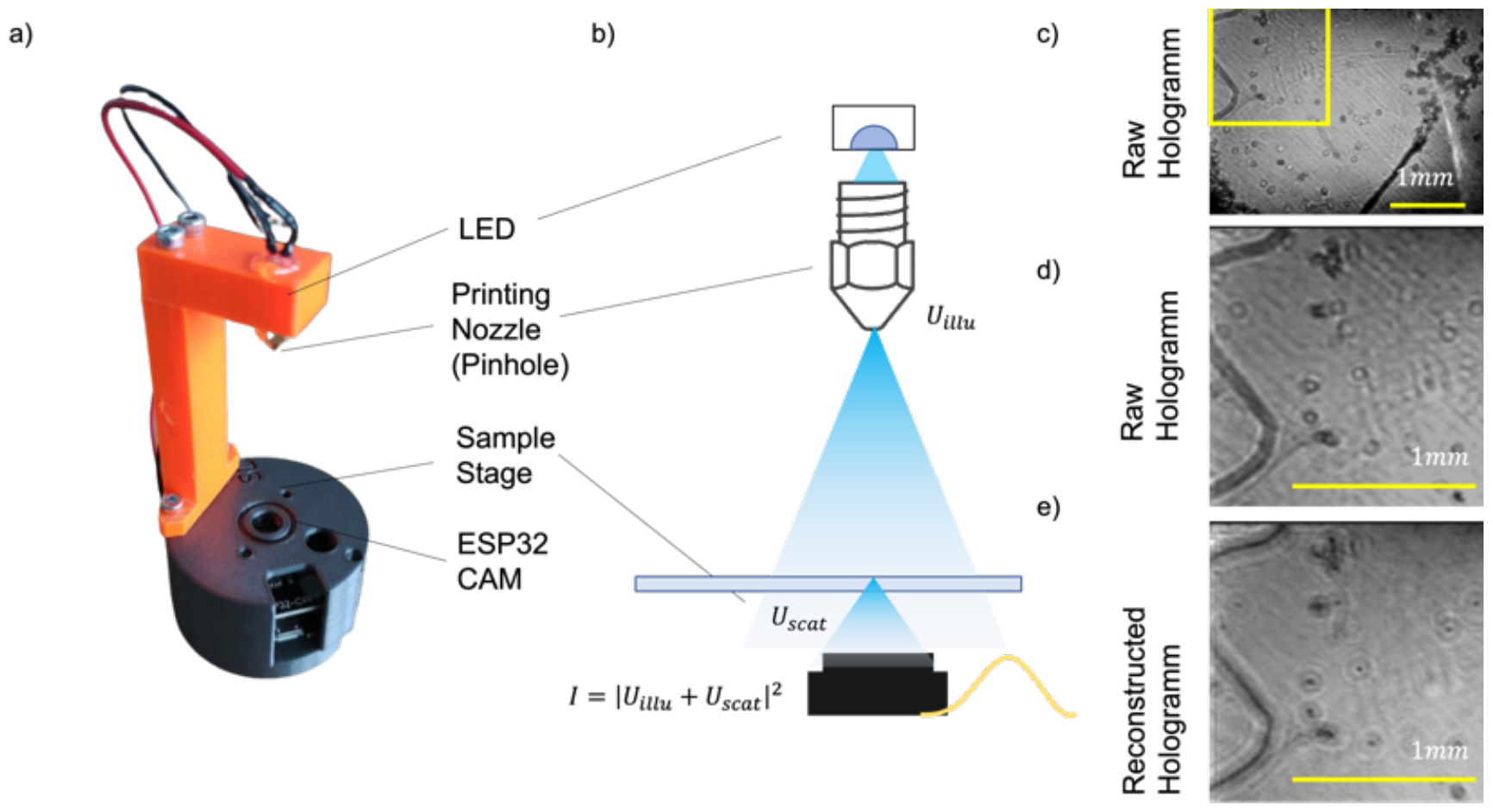
a) Inline holographic microscope, with an orange arm holding an LED whose output is spatially filtered by an FDM 3D printer’s hotend nozzle. b) The point source of light sends out spherical waves so that scattered and unscattered waves interfere on the sensor. Using the Fresnel transform, the raw digital hologram (c, d) can be numerically refocused as in e).

These results with our prototype HoloESP configuration demonstrate the preliminary feasibility of adapting the ESPressoscope into configurations which integrate complex real-time image processing algorithms.

### Firmware and user interface

An easy-to-use digital microscope should be operable with a user interface which is simple and intuitive and which doesn’t require complex installation procedures. Basic functionalities such as focusing, image acquisition, timelapse initiation, and configuration of camera and imaging parameters should be easily accessible through this interface, and this interface should be easy to operate with minimal software and hardware tooling. ESP32 microcontroller boards are compatible with the Arduino Integrated Development Environment (IDE) which offers a wide range of libraries useful for building firmware with a user interface that can be operated from the web browsers of devices with Wi-Fi connectivity, such as computers and smartphones. The use of a web browser-based user interface offers several advantages. First, it eliminates the need to install software on devices external to the microscope, making it more accessible to a wider range of users. Second, it allows for easy access to the microscope’s functionality from any device, regardless of its operating system or configuration.

In implementing the firmware for the various ESPressoscope configurations, we used available third-party libraries to achieve a fully functional prototype. Additionally, in order to avoid requiring users to install third-party software for compiling or flashing our firmware onto the ESP32 module, we used the experimental Web Serial API as part of a web page (https://matchboxscope.github.io/firmware/FLASH.html) which can communicate with an ESP32 camera module connected by a USB cable; this allows the web page to upload firmware to the module, but currently only recent versions of Chromium-based web browsers implement the Web Serial API. Despite the limited browser support, this zero-cost, zero-installation setup process is important for reducing the expertise required to bring up, operate, and maintain a scientific instrument meant to be widely accessible. The firmware binaries are automatically compiled on cloud infrastructure with a continuous integration pipeline running in Github Actions (Supplement Figure 6). Firmware binaries can be uploaded using either an over-the-air (OTA) mechanism or through the Web Serial-based web page, which provides a selection of customized firmware configurations (e.g. for the ESP32-CAM board, the Seeed Studio Xiao Sense board, Serial Camera, ESPectrophotometer).

The resulting software has the following functionalities, all of which are accessible through the browser-based GUI (Supplement Figure 6):

- Wi-Fi network and internet connectivity management
- Live preview from the camera for exploring samples and adjusting focus
- Capture and saving of images to the ESP32 camera board’s attached micro-SD card
- Timelapse imaging, where the microcontroller periodically captures images, saves them to an external micro-SD card, and otherwise stays in a deep sleep mode to reduce power consumption (Figure S1)
- Focus adjustment, where the lens can be moved to increase focusing contrast
- Configuration of camera and image acquisition parameters, such as exposure time, gain, resolution, and timelapse interval
- Control of hardware parameters specific to optional hardware modules, for microscopes with those modules
- Image processing, using the built-in integration with ImJoy and ImageJ.js [31] Uploading of new firmware using the Over-the-Air (OTA) update feature

All functionalities are implemented using an HTTP-based REST API and a Javascript-based frontend which is served by a web server running on the ESP32 camera board’s microcontroller. Alternatively, we provide a very basic Android app (https://matchboxscope.github.io/docs/APP) which uses the REST API and has the ability to save images from the microscope to the phone’s storage. This is especially helpful for acquiring live video clips from the microscope, as higher image acquisition can be achieved by downloading frames via HTTP and saving them from a computer than by having ESP32 camera module to save individual frames on its attached micro-SD card.

## Conclusions and Outlook

In this work, we aimed to develop a cost-effective microscopy system that could be adapted for many different settings and use-cases, including in situ deployments in isolated or otherwise challenging environments. Our described design, the ESPressoscope platform, enables people to make mission-specific trade-offs between various priorities - component availability, component cost, simplicity of assembly, provided functionalities, and imaging quality - by specifying a variety of recomposable modules which can be substituted for each other in the design of a specific microscope configuration. We have also introduced and preliminarily characterized some specific prototypes as easy-to-replicate ESPressoscope configurations which enable a variety of deployment scenarios. In particular, the Anglerfish configuration of the ESPressoscope platform is - to the best of our knowledge - the first low-cost underwater microscope which aims to be reproducible by a layperson and to be deployable in situ at larger scales. We believe that the kinds of observations Anglerfish enables for microbiota on aquatic surfaces in ecological contexts may provide new information about the biodiversity and behaviors of aquatic microbial communities.

The current implementations of our described ESPressoscope configurations have a number of limitations and problems to be solved in future work. For example, the ESP32-CAM module has been reported by some users to behave unexpectedly and to break for no obvious reason. This may be due to manufacturing mistakes, inappropriate selection of components such as the power controller, or design issues with the ESP32-CAM module itself. One known design issue of the ESP32-CAM board, for example, is inadequate isolation of the ground planes, leading to occasional noise on the board’s input pins. This lack of reliability may lead to unexpected errors in ESPressoscope firmware or even in the failure of a timeseries experiment. Because the ESP32-CAM board is just a composable module within the ESPressoscope architecture, any problems inherent to the board can be solved by substituting it with an alternative ESP32 camera board - potentially increasing the total cost of the microscope’s parts - as long as the substitute board can be integrated into the ESPressoscope’s mechanical structure with a 3D-printable adapter. We have already done this with the Seeed Studio Xiao Sense board for the Anglerfish configuration with UC2 modules. Similarly, the ESPressoscope’s modularity enables other iterative improvements to be made in the design of new microscope configurations to improve device reliability or other aspects of system performance.

The ESPressoscope configurations and modules described in this paper all emphasize low cost and ease of replication in their design tradeoffs, and thus they have limitations in image quality, compared to other open-source systems such as the OpenFlexure microscope [8]. Nevertheless, this work may still be useful for many microscopy-related projects in field research and in educational settings (exemplary shown in Figure 7), whether on their own or in combination with other microscopy systems. With the provided capabilities, low cost, and use of widely-available parts in the ESPressoscope configurations we have described, we believe these microscopes could become valuable image data collection tools for many communities. We plan to begin production scale-up with 3D-printing and low-volume injection molding using desktop injection molding machines.

**Fig 7.**
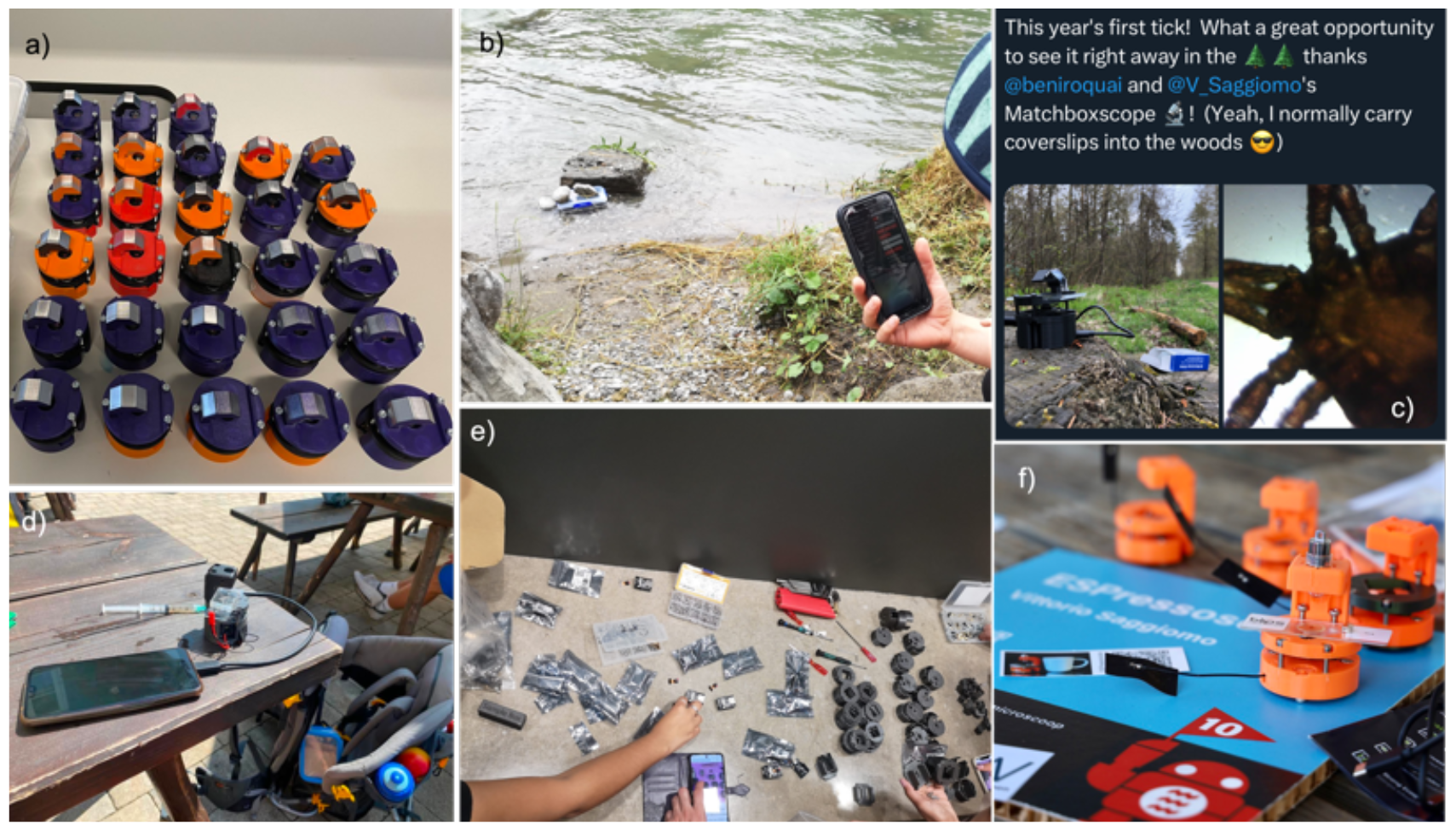
Figure 7 (a) The ESPressoscope can easily be mass-produced to deploy it in different settings such as field work (b, d) or STEM education (e). The light weight and small size of the minimal Matchboxscope configuration make it suitable in remote locations for on-site measurements, as already demonstrated by various users under the hashtag “#matchboxscope” on Twitter/X. (f) The recent Seeed Studio XIAO Sense ESP32 camera board makes the device even more compact, as presented at the 2023 Eindhoven Maker Faire.

The ESPressoscope’s design process followed a tight feedback loop with users whose task was to replicate the design only with the help of the provided online documentation during a series of workshops in multiple countries (e.g. Nigeria, Netherlands, Germany, USA UK). These workshops also helped us to improve the online documentation for users beyond those workshops. Ahead of this manuscript, users have already started posting on social media under the hashtag “#matchboxscope” about new microscopes being set up and new data being collected, which gives us an idea of who is using this microscope and why.

User feedback on social media as well as on GitHub and in a dedicated non-representative survey has improved the device during the still-ongoing open development process. It also demonstrates that pre-publication and active interaction with the community, rather than being a perceived stealing of ideas, instead facilitates improvements to projects, as users were involved early in the design process and experimentation. Thus, ESPressoscope contributes to research on how Open Science can be designed and conducted with the help of open instrumentation. Now, we invite the community to improve the system by giving feedback and reporting availability of ESPressoscope components described in this paper, in order to develop an even more capable and widely-available device.

## Acknowledgements

We thank Stephan Saalfeld from the HHMI Janelia Research Farm and the AQLM Course at the MBL in Woods Hole MA, USA for conducting a workshop on building a series of ESPressoscopes. We thank all users of the ESPpressoscope platform for valuable feedback which has improved the hardware and software prior to publication. We acknoledge funding from the Nexus GIF grant no. G-1566-143.13/2023. We thank the free state of Thuringia for supporting this project.

## Notes

https://matchboxscope.github.io/

### Competing Interest Statement

The authors have declared no competing interest.

## References

[1] M. Del et al. “The Field Guide to 3D Printing in Optical Microscopy for Life Sciences”. In: Adv. Biology 6 (2022), p. 2100994.

[2] J.W.P. Brown, A. Bauer, M.E. Polinkovsky, et al. “Single-molecule detection on a portable 3D-printed microscope”. In: Nat Commun 10 (2019), p. 5662.

[3] B. Diederich et al. “CellSTORM—Cost-effective super-resolution on a cellphone using dSTORM”. In: PLOS ONE 14.1 (2019), e0209827.

[4] Benedict Diederich et al. “Nanoscopy on the Chea(i)p”. In: bioRxiv (2020). eprint: https://www.biorxiv.org/content/early/2020/09/04/2020.09.04.283085.full.pdf. URL: https://www.biorxiv.org/content/early/2020/09/04/2020.09.04.283085.

[5] Ando C. Zehrer et al. “An open-source, high resolution, automated fluorescence microscope”. In: (2023).

[6] P. D. Gordon et al. “A portable brightfield and fluorescence microscope toward automated malarial parasitemia quantification in thin blood smears”. In: PLOS ONE 17.4 (2022), e0266441.

[7] Hongquan Li et al. “Octopi: Open configurable high-throughput imaging platform for infectious disease diagnosis in the field”. In: (2019).

[8] Joel T. Collins, Joe Knapper, Julian Stirling, et al. “Robotic microscopy for everyone: the OpenFlexure microscope”. In: Biomed. Opt. Express 11 (2020), pp. 2447–2460.

[9] Hongquan Li et al. “Squid: Simplifying Quantitative Imaging Platform Development and Deployment”. In: (2020).

[10] PJ Tadrous. “PUMA – An open source 3D printed direct vision microscope with augmented reality and spatial light modulator functions”. In: Journal of Microscopy 283 (2021), pp. 259–280.

[11] E.C. Orenstein et al. “The Scripps Plankton Camera system: A framework and platform for in situ microscopy”. In: Limnol Oceanogr Methods 18 (2020), pp. 681–695.

[12] R.W. Bowman. “Improving instrument reproducibility with open source hardware”. In: Nat Rev Methods Primers 3 (2023), p. 27.

[13] B. Diederich, C. Müllenbroich, N. Vladimirov, et al. “CAD we share? Publishing reproducible microscope hardware”. In: Nat Methods 19 (2022), pp. 1026–1030.

[14] J. Hohlbein, B. Diederich, B. Marsikova, et al. “Open microscopy in the life sciences: quo vadis?” In: Nat Methods 19 (2022), pp. 1020–1025.

[15] D. Zakoth et al. “Open Source Photonics at the Abbe School of Photonics: How Makerspaces foster Open Innovation Processes at Universities”. In: Fifteenth Conference on Education and Training in Optics and Photonics: ETOP 2019. Optica Publishing Group. 2019, pp. 11143–162.

[16] J. S. Cybulski, J. Clements, and M. Prakash. “Foldscope: Origami-Based Paper Microscope”. In: PLOS ONE 9.6 (2014), e98781.

[17] A.H. Velders et al. “Step-by-step: A microfluidic (PDMS) staircase device for size sorting microparticles down to 25μm using a 3D-printed mold”. In: ChemRxiv (2023). This content is a preprint and has not been peer-reviewed.

[18] Joel T. Collins et al. “Simplifying the OpenFlexure microscope software with the web of things”. In: R. Soc. open sci. 8211158211158 (2021).

[19] J. Baina and J. Dublet. “Automatic focus and iris control for video cameras”. In: (1995), pp. 232–235.

[20] World Health Organization. Schistosomiasis Fact Sheet. 2022. URL: https://www.who.int/news-room/fact-sheets/detail/schistosomiasis.

[21] Luisa Galgani et al. “Environmental Science & Technology”. In: 56.22 (2022), pp. 15638–15649.

[22] Christoph D. Rummel et al. “Impacts of Biofilm Formation on the Fate and Potential Effects of Microplastic in the Aquatic Environment”. In: Environmental Science & Technology Letters 4.7 (2017), pp. 258–267. URL: 10.1021/acs.estlett.7b00164.

[23] Aditya R. Nayak et al. “A review of holography in the aquatic sciences: in situ characterization of particles, plankton, and small scale biophysical interactions”. In: Frontiers in Marine Science 7 (2021), p. 572147.

[24] A. Mullen, T. Treibitz, P. Roberts, et al. “Underwater microscopy for in situ studies of benthic ecosystems”. In: Nat Commun 7 (2016), p. 12093.

[25] K. Shahani, H. Song, S.R. Mehdi, et al. “Design and Testing of an Underwater Microscope with Variable Objective Lens for the Study of Benthic Communities”. In: J. Marine. Sci. Appl. 20 (2021), pp. 170–178.

[26] J. Schindelin, I. Arganda-Carreras, E. Frise, et al. “Fiji: an open-source platform for biological-image analysis”. In: Nat Methods 9 (2012), pp. 676–682.

[27] Mini Peristaltic Pump for 28BYJ-48 Stepper Motor. Accessed 2024. URL: https://www.printables.com/model/63352-mini-peristaltic-pump-for-28byj-48-stepper-motor/collections.

[28] Thibaut Pollina et al. “PlanktoScope: Affordable Modular Quantitative Imaging Platform for Citizen Oceanography”. In: Frontiers in Marine Science 9 (2022). ISSN: 2296-7745. URL: https://www.frontiersin.org/articles/10.3389/fmars.2022.949428.

[29] Public Lab. Spectrometry. Accessed 2024. URL: https://publiclab.org/wiki/spectrometr

[30] GaudiLabs. 3DFiberSpectrograph. commit 9c3a729. 2023. URL: https://github.com/GaudiLabs/3DFiberSpectrograph.

[31] W. Ouyang, F. Mueller, M. Hjelmare, et al. “ImJoy: an open-source computational platform for the deep learning era”. In: Nat Methods 16 (2019), pp. 1199–1200.

